# Bayesian Inference for Brain Source Imaging with Joint Estimation of Structured Low-rank Noise

**DOI:** 10.1101/2023.03.19.533348

**Authors:** Sanjay Ghosh, Chang Cai, Yijing Gao, Ali Hashemi, Stefan Haufe, Kensuke Sekihara, Ashish Raj, Srikantan S. Nagarajan

**Affiliations:** University of California San Francisco, USA; Central China Normal University, China; Technical University of Berlin, Germany; Signal Analysis Inc., Japan

**Keywords:** EEG, MEG, brain source imaging, low-rank noise

## Abstract

The inverse problem in brain source imaging is the reconstruction of brain activity from non-invasive recordings of electroencephalography (EEG) and magnetoencephalography (MEG). One key challenge is the efficient recovery of sparse brain activity when the data is corrupted by structured noise that is low-rank noise. This is often the case when there are a few active sources of environmental noise and the MEG/EEG sensor noise is highly correlated. In this paper, we propose a novel robust empirical Bayesian framework which provides us a tractable algorithm for jointly estimating a low-rank noise covariance and brain source activity. Specifically, we use a factor analysis model for the structured noise, and infer a sparse set of variance parameters for source activity, while performing Variational Bayesian inference for the noise. One key aspect of this algorithm is that it does not require any additional baseline measurements to estimate the noise covariance from the sensor data. We perform exhaustive experiments on both simulated and real datasets. Our algorithm achieves superior performance as compared to several existing benchmark algorithms.

## 1. INTRODUCTION

Electromagnetic brain imaging is the reconstruction of brain activity from non-invasive recordings of magnetic fields and electric potentials. Electroencephalography (EEG) and magnetoencephalography (MEG) are widely used techniques to study the function of human brain [1, 2]. Efficient estimation of the brain activity on the cortex surface is important for neuroscience research and clinical diagnosis [3]. It is crucial to determine, to the extent possible, where and when neurophysiological activity is occurring. Thus, the inverse problem turns out to be estimating both the spatial location of the sources and the time course of brain activity from the EEG/MEG measurements. Several methods have been introduced to solve this ill-posed inverse problem of brain source imaging. The regularization based methods attempt to enforce prior information on the source signal. The minimum-norm estimation algorithm (MNE) [4] tries to minimize the *L*_2_ norm of the solution favoring low power of the brain activity. Other variants of MNE include the weighted MNE (wMNE) [5], low resolution brain electromagnetic tomography (LORETA) [6], and standardized LORETA (sLORETA) [7], etc. Another broad class of approach is to consider Bayesian techniques [8, 9, 10, 11, 12, 13, 14] with appropriate priors assigned to the model parameters. Recently introduced Champagne algorithm [8], a novel tomographic source reconstruction algorithm derived in an empirical Bayesian fashion with incorporation of deep theoretical ideas about sparse-source recovery from noisy, constrained measurements. One of active current research focus is to improve upon the reconstruction methods under realistic noise assumption such as low-rank statistical model [11]. An enduring challenge is to jointly remove the noise of the sensor arrays and recover the brain signal. The noise statistics in the model plays a crucial role in the success of sparse source recovery. In particular, the noise covariance in sensor data is what dictates the working of Bayesian frameworks. The existing works include noise covariance matrix of both diagonal and full structure. In this work, we consider one of most realistic assumptions - low rank noise covariance. This is often the case when there are a few active sources of environmental noise or the MEG/EEG sensors are highly correlated. To best of our knowledge, no existing algorithm has addressed the brain source estimation problem under low-rank noise covariance.

In this paper, we propose a novel robust empirical Bayesian framework for brain source imaging under low-rank noise assumption. It provides us a tractable algorithm for iteratively estimating the noise covariance and the brain source activity. The proposed algorithm is found to be quite robust to initialization and computationally efficient. The proposed algorithm does not require any additional baseline measurements to estimate noise covariance from sensor data. We further show that the new algorithm produces competitive performance with benchmark method on real MEG data and able to resolve distinct and functionally relevant brain areas.

This paper is outlined as follows. In Section 2, we introduce the inverse problem in brain source imaging under low-rank noise. This is followed by the proposed Bayesian reconstruction algorithm. Next, in Section 3, we show the experimental results of our approach on synthetic and real MEG data. Finally, we conclude in Section 4.

## 2. THEORY

### 2.1. Probabilistic Generative Model

In brain source imaging setup, brain activity originates from a number of electric current dipoles, where the location, orientation, and the magnitude of each dipole determine the recorded signal from the EEG electrodes. We consider the following forward model between measurements and brain sources:

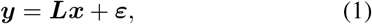

where ***y*** ∈ ℝ^*M ×K*^ is the EEG/MEG sensor data measured at *M* sensors at *K* time-points, ***x*** ∈ ℝ^*N ×K*^ is the underlined brain signals where *N* is the number of voxels. The lead-field matrix ***L*** ∈ ℝ^*M ×N*^ represents the propagation of electromagnetic field from a particular source location to the EEG/MEG sensors. The additive measurement noise ***ε*** is drawn from *N*(**0, Λ**^−1^), where **Λ** is the precision of the noise.

In this work, the noise is modelled in terms of the co-variance of the noise statistics as the follows:

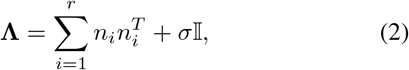

where 𝕀 ∈ ℝ^*M ×M*^ is an identity matrix, *σ* is a scaling factor, and *r* represents the rank of the noise statistics. The above low-rank noise is simulated by placing correlated passive (correlated) source of noise at close proximity of MEG/EEG sensors. Notice that for *σ* = 0, the noise co-variance **Λ** in either (2) is purely low-rank; it’s rank is *r*. For *σ* ≠ 0 in (2), the noise is a mixture of low-rank and homoscedastic statistics. We assume zero-mean Gaussian prior for the underlying source: *x*_*k*_ ∈ ℝ^*N ×*1^ ∼ 𝒩 (0, **Φ**^−1^), *k* = 1, …, *K*, where 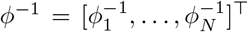 contains *N* distinct unknown variances for the brain sources and **Φ**^−1^ = diag(*ϕ*^−1^). We consider to solve the inverse problem by using Bayesian learning framework; by finding the maximum a-posterior probability (MAP) solution. The posterior probability *p*(*x*_*k*_|*y*_*k*_) can be derived by using Bayes’ rule

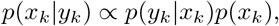

where *p*(*y*_*k*_|*x*_*k*_) = 𝒩 (***L****x*_*k*_, **Λ**^−1^) and *p*(*x*_*k*_) = 𝒩 (*x*_*k*_|0, **Φ**^−1^). It is straightforward to show that posterior probability *p*(*x*_*k*_|*y*_*k*_) is also Gaussian. Suppose, the posterior probability takes the following form:

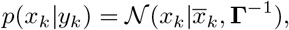

where 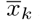 is the posterior mean, and **Γ** is the posterior precision matrix. Further it can be shown that:

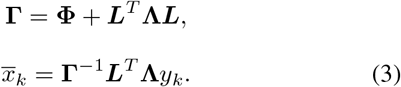

### 2.2. Proposed Bayesian Inference Algorithm

Notice that we need both **Φ** and **Λ** to compute the Bayesian estimate of unknown *x* is computed using (3). However, since we have access to only ***y***, the idea is to compute the complete data likelihood and average over it with the posterior probability as follows:

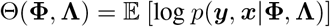

where the expectation [·] is taken with respect to *p*(***x***|***y***). The complete data likelihood is expressed as

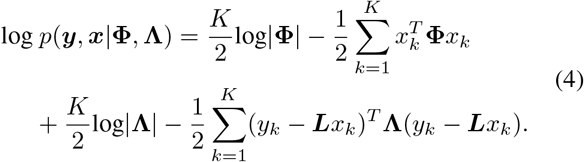

#### Update of Source Parameter

Here we obtain the optimal 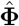 by maximizing the average likelihood Θ(**Φ, Λ**) for a fixed **Λ** as follows: 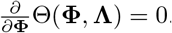. With further derivation, we can show that it follows MacKay update equation:

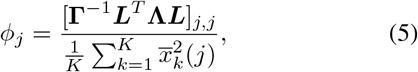

where [·] indicates (j, j)-th element of a matrix.

#### Update of Noise Parameter

In this work, we propose to estimate 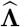 via factor analysis of the following residual noise at l-th iteration:

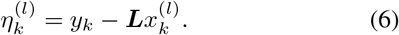

The idea is to obtain the covariance of 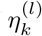 using Bayesian factor analysis. In other words, we assume a factored decomposition of *η*_*k*_ as follows:

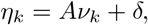

where the residual noise at *k*-th time-point is *η*_*k*_ ∈ ℝ^*M ×*1^, *A* ∈ ℝ^*M ×P*^, *ν*_*k*_ is a *P* -dimensional column vector, and *δ* is modeling noise. Notice that we drop the iteration symbol *l* for simplification of notations.

##### Algorithm 1: Low-rank Champagne

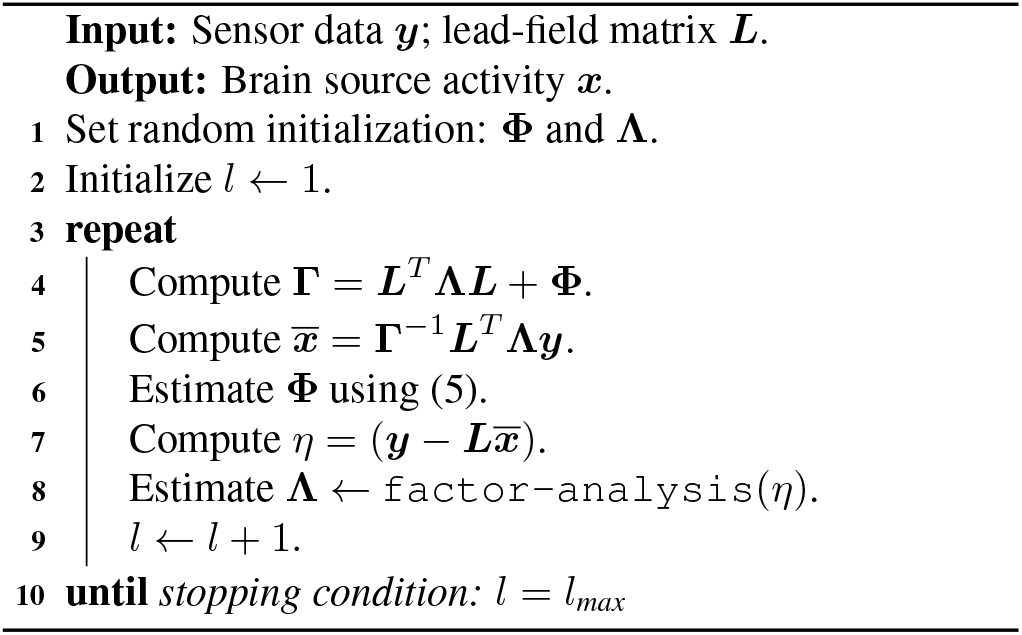

We assume the prior probability distribution of the factor ***ν***_*k*_ is assumed to be the zero-mean Gaussian with its precision matrix equal to the identity matrix,

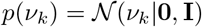

The factor activity is assumed to be independent across time. Thus, the joint prior distribution:

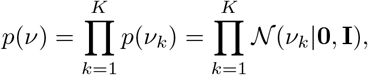

where **I** is an identity matrix of size (*P×P*). The modeling noise *δ* is assumed to be Gaussian with the mean of zero:

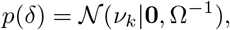

where Ω is a diagonal precision matrix. Further, with few steps of mathematical derivations, we obtain the Bayesian estimate of residual noise covariance as follows:

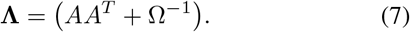

Finally we summarize the proposed reconstruction method in Algorithm 1. The key aspect is the novel way of estimating covariance of residual noise within each iteration of the proposed Bayesian algorithm. We note that our work is motivated by one recent empirical Bayesian source reconstruction algorithm is called Champagne [8]. We refer the proposed algorithm as Low-rank Champagne.

## 3. RESULTS

### 3.1. Simulations and Real Data

We generate source signal data by simulating dipole sources with fixed orientation. The damped sinusoidal time courses with frequencies sampled randomly between 1 and 75 Hz are created as voxel source time activity. The time-courses are then projected to the sensors using the lead-field matrix generated by the forward model. We consider 271 MEG sensors and a single shell spherical model [15] implemented in SPM12 http://www.fil.ion.ucl.ac.uk/spm at the default spatial resolution of 8196 voxels at approximately 5 mm spacing. We set time period as 480 samples with source activities of interest and noise activity.

Real MEG data we used in our experiments was acquired in the Biomagnetic Imaging Laboratory at University of California, San Francisco (UCSF) using a CTF Omega 2000 whole-head MEG system from VSM MedTech (Coquitlam, BC, Canada) with 1200 Hz sampling rate. The lead field for each subject was calculated in NUTMEG software using a single-sphere head model (two spherical orientation lead fields) and an 8 mm voxel grid. the lead field was calculated using a three-shell spherical model at the coarse resolution.

### 3.2. Benchmarks and Performance Quantification

In this paper, we choose noise-learned Champagne [Cai et al. 2021] as the key benchmarks to compare with. Two other we compare are sLORETA [7] and full structure noise (FUN) learning method [16]. The performance of simulated brain source reconstruction is evaluated based on response receiver operator characteristics (FROC) [12]. We compute *A*^*′*^ and aggregated performance (AP) metrics to quantify performance of source localization and reconstruction. For both, a higher value indicates better result.

### 3.3. Simulation Results

A brain source reconstruction example is shown in Fig. 1. We simulated a time-course with 5 active sources randomly placed with 3D brain. We also generated low-rank noise as demonstrated in (2). The simulated brain signal is then projected to measurement space by applying the leadfield matrix and then the noise is added at SNR = 5dB. Finally, we reconstruct the underlined time-course using the proposed algorithm. We also compared the reconstruction performance with recent method by Cai et al. [12]. It is visually evident that our method performs better that the state-of-the-art method.

**Fig. 1.**
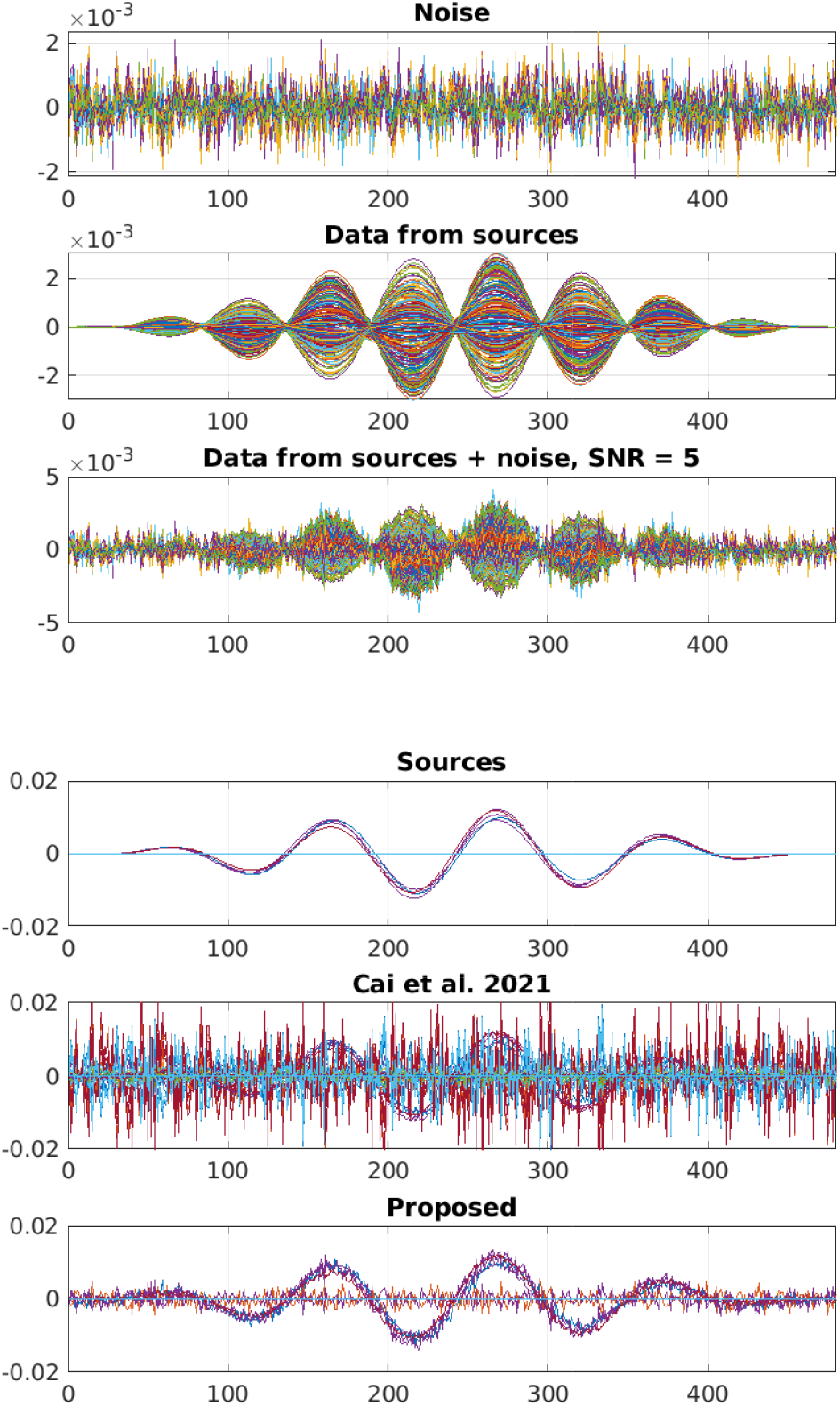
Reconstruction of active brain sources under simulated brain noise of rank *r* = 5 and SNR 5 dB.

Performance results versus SNR (for a fixed rank) for the respective algorithms are plotted in Fig. 3. Reconstruction performance is evaluated for five randomly seeded dipolar sources with an inter-source correlation coefficient of 0.99. We also study the performance with varying the rank of the noise covariance for fixed SNR level in Fig. 4. It is clear that proposed *Low-rank Champagne* outperforms in both setting in terms of both *A*^*′*^ and AP metrics.

**Fig. 2.**
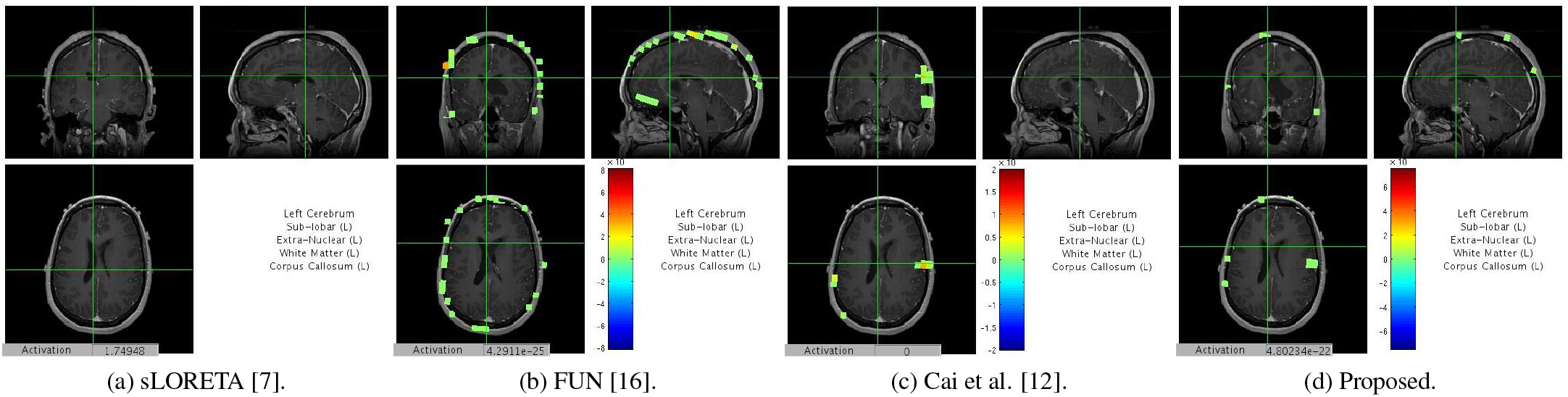
Auditory evoked field (AEF) results.

**Fig. 3.**
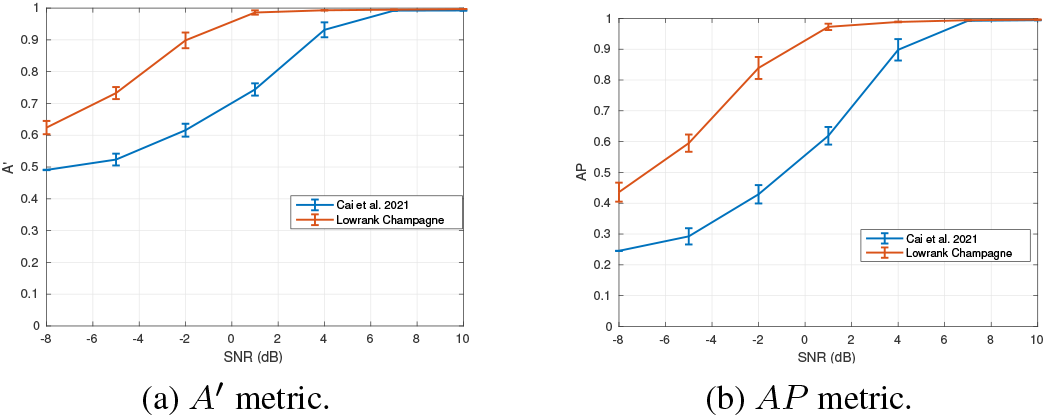
Performance with varying SNR (dB) for rank r = 10.

**Fig. 4.**
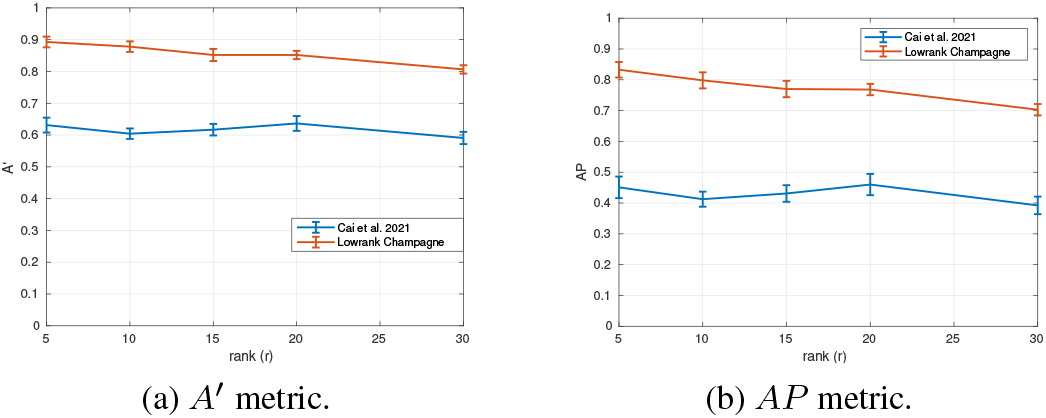
Performance with varying rank for -2 dB SNR.

### 3.4. Results on real datasets

The localization result for AEF (Auditory Evoked Fields) data of a single representative is shown in Fig 2. We also compare the performance with sLORETA, FUN, and Cai et al. [12]. Our method is able to localize bilateral auditory activity which is supposed to be in Heschl’s gyrus, the location of primary auditory cortex. In contrast, sLORETA performance is not as good because of the presence of correlated sources.

## 4. DISCUSSION

In this paper, we proposed a new method for MEG source reconstruction and localization. The algorithm is able to estimate sensor noise from observed data without the need for additional pre-stimulus or baseline data. To be the best of our understanding, the improved performance of this algorithm arises from the efficient the method of estimating the noise statistics via factor analysis of residual component. In summary, this algorithm displays significant advantages over many existing benchmark algorithms for electromagnetic brain source imaging.

